# Probabilistic estimation of identity by descent segment endpoints and detection of recent selection

**DOI:** 10.1101/2020.07.15.205179

**Authors:** Sharon R. Browning, Brian L. Browning

## Abstract

Most methods for fast detection of identity by descent (IBD) segments report identity by state segments without any quantification of the uncertainty in the endpoints and lengths of the IBD segments. We present a method for determining the posterior probability distribution of IBD segment endpoints. Our approach accounts for genotype errors, recent mutations, and gene conversions which disrupt DNA sequence identity within IBD segments. We find that our method’s estimates of uncertainty are well calibrated for homogeneous samples. We quantify endpoint uncertainty for 7.7 billion IBD segments from 408,883 individuals of White British ancestry in the UK Biobank, and use these IBD segments to find regions showing evidence of recent natural selection. We show that many spurious selection signals are eliminated by the use of unbiased estimates of IBD segment endpoints and a pedigree-based genetic map. Nine of the top ten regions with the greatest evidence for recent selection in our scan have been identified as selected in previous analyses using different approaches. Our computationally efficient method for quantifying IBD segment endpoint uncertainty is implemented in the open source ibd-ends software package.

## Introduction

Pairs of individuals within a population can share one or more long segments of their genomes identical by descent due to inheritance from common ancestors. Identity by descent (IBD) segments are used in many applications, such as estimation of kinship,^1–3^ recent demography,^4–8^ mutation rates,^9–12^ and recombination rates,^13^ and detection of recent selection.^14–16^

The three main types of test for recent positive selection are based on population differentiation, admixture proportions, and haplotype structure. The first type looks for variants that differ markedly in frequency between populations.^17; 18^ The second type looks for regions in which the sample ancestry proportions in admixed individuals differ from those elsewhere in the genome.^19; 20^ One subtype of this test involves archaic admixture, such as introgression from Neanderthals into modern humans, and searches for regions in which the frequency of the archaic haplotype in a modern population is unusually high.^21^ The third type looks for high-frequency haplotypes that are unusually long.^22; 23^ IBD-based selection scans fall into this category.^14^ IBD scans look for genomic regions that have a significantly higher than average number of IBD segments. If the genome were completely neutral, and there are no biases in detecting IBD segments or estimating their centiMorgan (cM) lengths, the expected number of IBD segments exceeding some cM length threshold would be constant across the genome. In contrast, if certain haplotypes in a genomic region have a selective advantage, the effective size of the population is reduced in that region, which leads to a higher than expected number of IBD segments. IBD-based tests can also detect the effects of negative selection and balancing selection, since any type of selection will tend to decrease the effective population size within the genomic region.

An IBD segment for a pair of haplotypes is a segment of DNA inherited from a single common ancestor, with no crossovers occurring within the segment in the lineages of the two haplotypes since the common ancestor.^4; 6^ Within a shared IBD segment, sequence identity can be disrupted by mutation and gene conversion. In addition, genotype error can cause two haplotypes to appear to be discordant at a position. At such positions, two “identical by descent” haplotypes are in fact not identical. This non-identity needs to be considered when detecting IBD segments.

In the human genome, de novo single nucleotide mutations occur at an average rate of around 1.3 × 10^−8^ per base pair per meiosis,^12^ which is similar to the average rate of crossing over per base pair per meiosis. Thus, regardless of the number of generations since the most recent common ancestor, an average of approximately one mutation is expected in the lineage of an IBD segment.

For a pair of haplotypes drawn at random from a population, most of the genome is comprised of very short segments of IBD, with a very large number of generations to the most recent common ancestor. Since each short segment contains an average of approximately one discordance caused by mutation in addition to discordances caused by gene conversion, a series of closely-spaced discordances is a clear indication that the genomic interval containing the discordances is comprised of a sequence of short IBD segments. In contrast, when one observes a long segment without discordances, it is usually (depending on the population’s demographic history) highly probable that this segment is primarily comprised of a single long IBD segment resulting from recent common ancestry.

There are three primary paradigms for IBD segment detection. The first paradigm considers a pair of haplotypes to be either “IBD” or “not IBD” at each position in the genome. A hidden Markov model, with pre-determined IBD proportion and rates of transition between the IBD and non-IBD states, may be used to obtain posterior probabilities of IBD and non-IBD at each position.^24–29^ This paradigm developed out of the analysis of pedigree data, and is very natural in that setting.^30^ However, for population data with unknown relationships, the dichotomy into IBD and non-IBD is artificial and ignores the fact that each pair of haplotypes has a common ancestor at each position in the genome, although that ancestor may have lived a long time ago.

The second paradigm considers the length of shared segments. A segment is identical by descent if it is inherited from a common ancestor and exceeds a length threshold. In practice, if identity by state (IBS) sharing extends beyond some threshold, the segment is reported as identical by descent.^31–33^ This paradigm recognizes the potential existence of IBD segments that are shorter than the threshold, but does not try to find them. The threshold is typically chosen to be a length above which the accuracy of the reported segments is high.^31; 32^

The third paradigm considers two haplotypes to be identical by descent if their time to most common ancestor (TMRCA) is less than some specified number of generations.^16; 34^

In this work we take a different perspective. We recognize that a pair of haplotypes is, strictly speaking, identical by descent at every point in the genome. However, for any given point in the genome, the endpoints of the IBD segment containing that point are unknown (Figure 1A). It is possible that one or more discordances at the end of the segment are actually contained within the long IBD segment (Figure 1B). It is also possible that IBD ends before IBS ends, so that the end of the IBS segment is not part of the long IBD segment, but instead contains one or more neighboring short IBD segments (Figure 1C). In some cases, two or more long IBD segments in a region can be mistaken for a single long IBD segment (Figure 1D).^35^ Our approach quantifies this uncertainty. In Results, we show that our quantification is well-calibrated, and we apply our method to perform an IBD-based selection scan in the UK Biobank.

**Figure 1:**
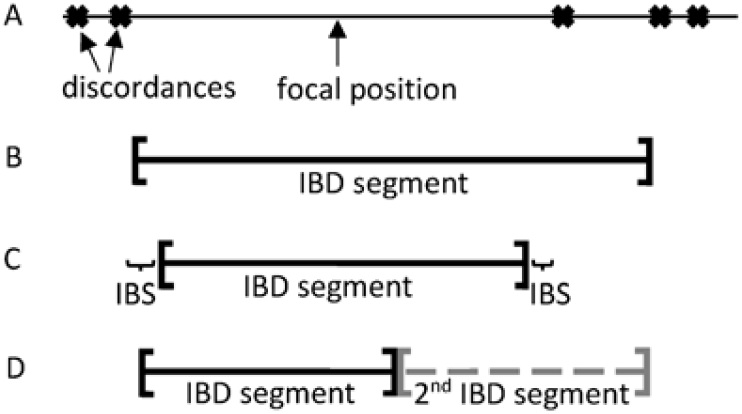
Uncertainty in IBD endpoints. A: Allele discordances between two haplotypes are represented as crosses. We wish to estimate the endpoints of the IBD segment that covers the focal position in the middle of the longest identity by state (IBS) interval. B-D: Three of the possibilities for the shared IBD segment that covers the focal position. B: The IBD segment contains the first discordance to the right of the focal position. C: The IBD segment does not extend all the way to the discordances and has short flanking segments of IBS. D: Two moderately long IBD segments are adjacent. In this case, the second IBD segment is not of direct interest because it does not cover the focal position.

## Methods

### Overview of method

The input data for our method are phased genotypes and candidate IBD segments. Highly-accurate phased genotypes can be obtained from statistical phasing in large cohorts of accurately-genotyped individuals.^36^ The candidate IBD segments may be obtained using a length-based IBD detection method such as hap-ibd.^33^ Our method estimates the posterior probability distributions of the endpoints of the candidate IBD segments and outputs quantiles and samples from these posterior distributions.

Before estimating segment endpoints, we apply a minor allele frequency (MAF) filter to the phased genotypes that excludes variants with frequency less than 0.1%. In Results we show that these rare variants are not modelled as well as the more common variants and that including these rare variants negatively impacts the accuracy of the endpoint estimates.

We model allele discordance within an IBD segment using a user-specified error rate. Analysis results are not overly sensitive to the exact choice of error rate (see Results). Discordances within a segment are assumed to occur independently except when two or more closely-spaced discordances could have originated from the same gene conversion event.

We also model IBS extending beyond the end of an IBD segment. IBS segments can be comprised of multiple IBD segments. We do not try to directly model each of these IBD segments individually, but instead model the distribution of IBS segments found in the data. Short regions of IBS are modelled using the local context, because the IBS length distribution varies across the genome due to factors such as mutation rate and selection. Longer segments of IBS are modelled using chromosome-wide data because there is limited information about longer IBS segments from the local context.

We estimate the probability of the observed discordance data as a function of the IBD endpoints. We then use Bayes’ rule to obtain the probability distribution of each IBD endpoint. We work from a focal position within an IBS segment (Figure 1A) and estimate the probability distributions for the positions of the left and right endpoints of the IBD segment that covers the focal position.

### Notation

We wish to estimate the endpoints of the IBD segment covering a position *x*_0_ for a given pair of haplotypes, *H*_1_ and *H*_2_. All positions are measured in terms of genetic distance in Morgans, and haplotype phase is assumed to be known. In this description we are only concerned with the estimation of the right endpoint of the IBD segment covering *x*_0_. The estimation of left endpoints is similar. Index the markers to the right of *x*_0_ by 1, 2, 3,…, *M*, where *M* is the last marker on the chromosome. Let the positions of these markers be *x*_1_, *x*_2_, *x*_3_,…, *x_M_*.

The IBS data, *D*, is the observed IBS status (identical or discordant) for the alleles on haplotypes *H*_1_ and *H*_2_ at the markers to the right of *x*_0_. Let *D*[*a, b*] denote the IBS status at markers with indices *a* ≤ *i* ≤ *b*. Let *ϵ* be the average proportion of discordant markers within IBD segments (the “error rate” mentioned above). We approximate (1 − *ϵ*) with 1 and thus omit terms of (1 − *ϵ*) in our calculations.

### Modelling the IBS data for the IBD segment

We model the IBS data, *D*, from the focal point *x*_0_ rightwards as being generated by two processes. The first, up to the IBD endpoint *R*, requires that alleles should be identical except at a small number of discordances due to mutation, gene conversion, or genotype error. If discordances in the IBD segment are independent and the right endpoint is in the interval (*x_i_*, *x*_*i*+1_), the probability of the data in the part of the IBD segment to the right of the focal point (i.e. *P*(*D*[1, *i*])) is *ϵ^n_i_^*, where *n_i_* is the number of discordances between the first marker after the focal point and the *i*-th marker (inclusive) and factors of (1 − *ϵ*) are approximated by 1.

We use Bayes rule to obtain the posterior distribution of *R*, the position of the right endpoint (*R* > *x*_0_). For each inter-marker interval (*x_i_*, *x*_*i*+1_), *i* = 0,1,…, *M* − 1, the probability that the right endpoint is contained in the open interval (*x_i_*, *x*_*i*+1_) satisfies:

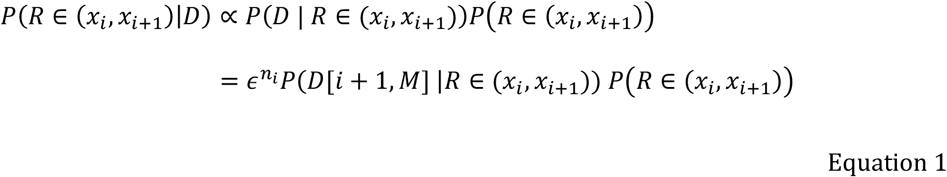

where “∝” denotes proportionality. Normalizing the probabilities in Equation 1 to sum to 1 over the *i* gives the posterior probability that the endpoint occurs in each interval (*x_i_*, *x*_*i*+1_).

The probability *P*[*R* ∈ (*x_i_*, *x*_*i*+1_)) is a prior probability for the position of the right endpoint, given the position *L*_0_ of the left endpoint (see “Iterative updating of endpoints and focal point” below). We model the population as having constant effective size 10,000 to obtain these probabilities. Details are given in Appendix 1.

The remaining component in Equation 1 is the probability *P*(*D*[*i* + 1, *M*] |*R* ∈ (*x_i_*, *x*_*i*+1_)), which is the probability of the IBS data to the right of the right IBD endpoint. We model this by considering the points of discordance as being the points of renewal in a renewal process. That is, we obtain probabilities of the length of each segment that contains all of the non-discordant positions up to and including the next discordance, with each such segment being treated as being independent. The probabilities of each of these sequences is obtained empirically from the observed data. Details are given in Appendix 2.

Appendix 3 describes how to obtain the posterior cumulative distribution function for the endpoint from the interval probabilities given in Equation 1.

### Iterative updating of endpoints and focal point

In the preceding section, we assumed that the left endpoint *L*_0_ was known, however it also needs to be estimated, and we do this estimation iteratively. We start by using the left endpoint of the input candidate IBD segment as the value of *L*_0_. After estimating the posterior distribution of the right endpoint, we use this distribution to obtain a new “right endpoint” *R*_0_ that is set equal to the 5^th^ percentile of this distribution. Percentiles are referenced by distance from the focal point *x*_0_. Thus, small percentiles are located closer to the focal point than larger percentiles. This choice of percentile is conservative (it reduces the estimated length of the IBD segment for the purpose of these calculations) and thus is likely to speed the convergence of the iterative approach. After estimating the right endpoint distribution, we estimate the left endpoint using the newly calculated value of *R*_0_ and obtain a new value of *L*_0_ as the 5^th^ percentile of the left endpoint distribution. Then if *L*_0_ − *x*_0_ is significantly altered (>10% change in length) from the previous value, we use the new value of *L*_0_ to re-estimate the right endpoint distribution and obtain a new value of *R*_0_. If *R*_0_ is significantly altered from the previous value, we use the new value of *R*_0_ to re-estimate the left endpoint, and so on. Whenever we change the value of *L*_0_ or *R*_0_ we update the focal point *x*_0_ which is located half-way between *L*_0_ and *R*_0_ in base coordinates. We perform a maximum of 10 updates of each endpoint. In order to prevent the focal point from moving outside the input candidate IBD segment, we constrain *L*_0_ and *R*_0_ to stay within the input candidate IBD segment.

### Estimation of error rate

After running the endpoint estimation algorithm, our ibd-ends software estimates the error parameter *ϵ* by measuring the rate of discordant alleles within inferred IBD segments. The procedure is as follows. For each IBD segment that has been analyzed with the endpoint estimation algorithm, take the interval bounded by the posterior 5^th^ percentile of the left and right endpoint distributions, and use these to obtain a cM length. If this length is < 2 cM, ignore the segment. Within the genomic region bounded by these endpoints, examine the alleles on the two IBD haplotypes. Count the number of mismatches, and the total number of positions examined. Across all segments, report the total number of mismatches, divided by the sum of the number of positions examined in each segment. If the estimated error rate does not differ significantly from the error rate used in the analysis (e.g. less than a three-fold difference), it is not necessary to re-run the analysis with the new value (see Results). For a large study, a pilot analysis on a small chromosome can be used to determine the error rate that should be used in the full analysis.

### Modified error rate to account for gene conversion

Gene conversions copy material from one haplotype to the other during meiosis and can thus result in discordant alleles between IBD haplotypes. The typical length of a gene conversion tract is around 300 base pairs.^39^ Changes will only occur at positions at which the individual in whom the gene conversion occurred was heterozygous. Thus, many gene conversions have no effect on allele discordance, but some gene conversions can result in more than one allele discordance occurring in proximity. Since these discordances are not independent events, we do not include an error term *ϵ* for each one, since that would be overly harsh and tend to result in premature truncation of the IBD segment. Instead, when more than one discordance occurs within 1 kb, we apply the error rate *ϵ* for the first discordance, and a less severe gene conversion error rate of *ϵ’* for each successive discordance within 1 kb of the first discordance (by default, *ϵ* = 0.0005 and *ϵ’* = 0.1).

### Analysis pipeline

Our software, ibd-ends, requires the input of candidate segments for which endpoints will be evaluated. In this work, we use hap-ibd^33^ to find the candidate segments. For many applications, one wishes to assess endpoint uncertainty for all IBD segments that exceed a length threshold. In that case, the key consideration is to avoid false negatives when detecting candidate IBD segments. If a potential IBD segment is not included in the input data to ibd-ends, it will not be included in the results. False positives (candidates for which the true IBD segment is actually shorter than the threshold) are less serious – they increase compute times but will be shown to be unlikely to be true long IBD segments when the segment endpoints are estimated. Thus, one should try to cast a wide net when identifying candidate IBD segments.

When analyzing sequence data, the high density of variants and the presence of genotype error can cause a high rate of discordances between IBD haplotypes. The hap-ibd method permits some discordances in a segment. It does so by finding seed IBS segments that exceed a certain length, and then extending these segments if there is another IBS segment that exceeds a minimum extension length and that is separated from the seed segment by a short non-IBS gap. One way to apply hap-ibd to sequence data is to reduce the seed and extension lengths, which effectively increases the permitted density of discordances, but this can significantly increase computation time. Here we take a different approach. We reduce the marker density of the sequence data for the hap-ibd analysis (but not for the ibd-ends analysis). We choose a minimum MAF for hap-ibd that reduces the marker density to approximately that of a 600k SNP array, and we apply this threshold using hap-ibd’s “min-mac” parameter. We retain the highest MAF variants because these are the most informative for detecting the candidate segments. This approach greatly reduces the density of variants, and thus reduces the number of IBD segments that would otherwise go undetected due to genotype error.

Except as otherwise noted, genotype data were phased using Beagle 5.1,^40^ input IBD segments for ibd-ends were obtained using hap-ibd,^33^ and default parameters were used for all programs.

### Simulation overview

We generated three sets of simulated data to investigate three conditions under which estimation of endpoints could be challenging: gaps in marker coverage, non-constant population size, and heterogeneous samples.

We added genotype error to the simulated data for each marker at a rate equal to the minimum of 0.02% and one-half the MAF for the marker. This error rate produces a discordance rate of 0.04% in markers with MAF > 0.04%, which matches the discordance rate seen in the TOPMed data^41^ and is six times higher than the 0.0067% discordance rate seen in the UK Biobank data.^42^ We also wanted to confirm that the method produces accurate results with higher error rates, so we added error to one data set at a rate equal to the minimum of 0.1% and one-half the MAF for the marker.

### Simulation of constant size population with variable marker density

We generated 60 Mb of data for 2000 individuals from a population with constant size of 10,000 diploid individuals. The recombination rate is 1 × 10^−8^ per base pair per meiosis. During the most recent 5000 generations, gene conversion initiations occurred at a rate of 2 × 10^−8^ and gene conversion tracts had a mean length of 300 base pairs. We used SLiM v3.3 to simulate the past 5000 generations,^43^ and msprime to add mutations and simulate the more distant past.^44; 45^ The mutation rate varied along the simulated region, with a new mutation rate each 100 kb that was uniformly distributed between 0 and 3 × 10^−8^ per bp per meiosis. In addition, we made a 1.5 Mb gap by removing genetic markers between positions 20.0 and 21.5 Mb, to represent a centromeric region. We added genotype error at a rate of 0.02% as described above. We used the simulated ancestral recombination graph to determine the true endpoints of IBD segments of length > 1 cM for all pairs of individuals within a subset of 500 individuals so that we could evaluate the accuracy of the inferred IBD segment endpoints.

We used the true (simulated) haplotypes, including any alleles changed by the addition of genotype error, for all analyses of these data. When detecting candidate IBD segments with hap-ibd, we used a minor allele count threshold of 1700 (minor allele frequency of 0.425; see Analysis pipeline), resulting in 10,760 markers after excluding markers in the 1.5 MB gap region, which corresponds to a mean density of one marker per 5.4 kb in the remaining 58.5 Mb. All markers with MAF > 0.1% (241,010 markers) were used in the ibd-ends analysis.

### Simulation of non-constant size population

We generated 60 Mb of data for 50,000 individuals from a UK-like population. These simulated data have been described previously.^33^ The demographic model has a population size of 24,000 in the distant past, a reduction to 3,000 occurring 5,000 generations ago, growth at rate 1.4% per generation starting 300 generations ago, and growth at rate 25% beginning 10 generations ago. The mutation rate is 1.3 × 10^−8^ per base pair per meiosis, while the recombination rate is 1 × 10^−8^ per base pair per meiosis. During the most recent 5000 generations gene conversion initiations occurred at a rate of 2 × 10^−8^ and gene conversion tracts had a mean length of 300 base pairs. We used SLiM v3.3 to simulate the most recent 5000 generations^43^ and msprime to add mutations and simulate the more distant past.^44; 45^ We generated two copies of the data: one with 0.02% added genotype error and one with 0.1% added genotype error as described above. We also created a SNP-array version of the data with 0.02% genotype error and 10,000 randomly selected markers with minor allele frequency > 5% (1 marker per 6 kb on average, corresponding to approximately 500k markers genome-wide). We used the simulated ancestral recombination graph to determine the true endpoints of IBD segments of length > 1 cM for all pairs of individuals within a subset of 1000 individuals so that we could evaluate the accuracy of the inferred IBD segment endpoints.

When applying hap-ibd to the UK-like sequence data we applied a minor allele count threshold of 4500 (minor allele frequency threshold of 0.45; see Analysis pipeline), resulting in 11,524 markers across the 60 Mb with a mean density of one marker per 5.2 kb. All markers with MAF > 0.1% (198,566 markers) were used in the ibd-ends analysis of the sequence data. When applying hap-ibd to the SNP-array data, we used the default minor allele count threshold (min-mac=2), and we analyzed all variants with frequency > 0.1% in the ibd-ends analysis of the SNP-array data.

### Simulation of a heterogeneous population

We simulated 10 Mb of data for 500 individuals of African-like ancestry and 500 individuals of European-like ancestry. The demographic history is the two-population model of Tennessen et al.,^46; 47^ implemented in stdpopsim.^48^ The combined sample represents an ancestrally heterogeneous population, which violates an assumption of our modelling of endpoint uncertainty. The recombination rate and mutation rate are both 1 × 10^−8^ per base pair per meiosis. We did not include gene conversion in the simulation. We simulated the data with msprime,^44^ and we added genotype error at a rate of 0.02% as described above. We used the simulated ancestral recombination graph to determine the true endpoints of IBD segments of length > 0.5 cM for all pairs of individuals so that we could evaluate the accuracy of the inferred IBD segment endpoints.

When applying hap-ibd to these data we applied a minor allele count threshold of 700 (MAF threshold of 0.35; see Analysis pipeline), resulting in 2215 markers across the 10 Mb with a mean density of one marker per 4.5 kb. We set the hap-IBD min-seed and min-output parameters to 1 cM since there are not many IBD segments of length > 2 cM in these data. All markers with MAF > 0.1% were used in the ibd-ends analyses (48,074 markers).

### UK Biobank data

We phased QC-filtered UK Biobank data (487,373 individuals) using Beagle 5.1,^40^ and then used hap-ibd to find candidate IBD segments among 408,883 White British individuals identified by the UK Biobank.^42^ We ran ibd-ends with default settings on the candidate IBD segments from the White British individuals to estimate the uncertainty in the endpoints of these IBD segments. We used Bherer et al.’s European genetic map which is based on family data from Iceland and other European populations.^49^ Variants located outside the bounds of the map are excluded from the ibd-ends analyses because extrapolated cM positions for markers outside the map can differ significantly from their true cM positions, leading to substantial under- or over-estimation of IBD segment lengths.

## Results

### Simulated data with variable marker density

Figure 2 shows the results of ibd-ends analysis on the simulated sequence data with constant population size, variable marker density, and a 1.5 Mb gap in marker coverage (see Methods for details). Even with the large gap and uneven marker density, the endpoints uncertainty is well-calibrated (Figure 2A), and coverage of sampled IBD segments across the simulated region is even (Figure 2B). 58% of sampled endpoints are located within 5 kb of the true value.

**Figure 2:**
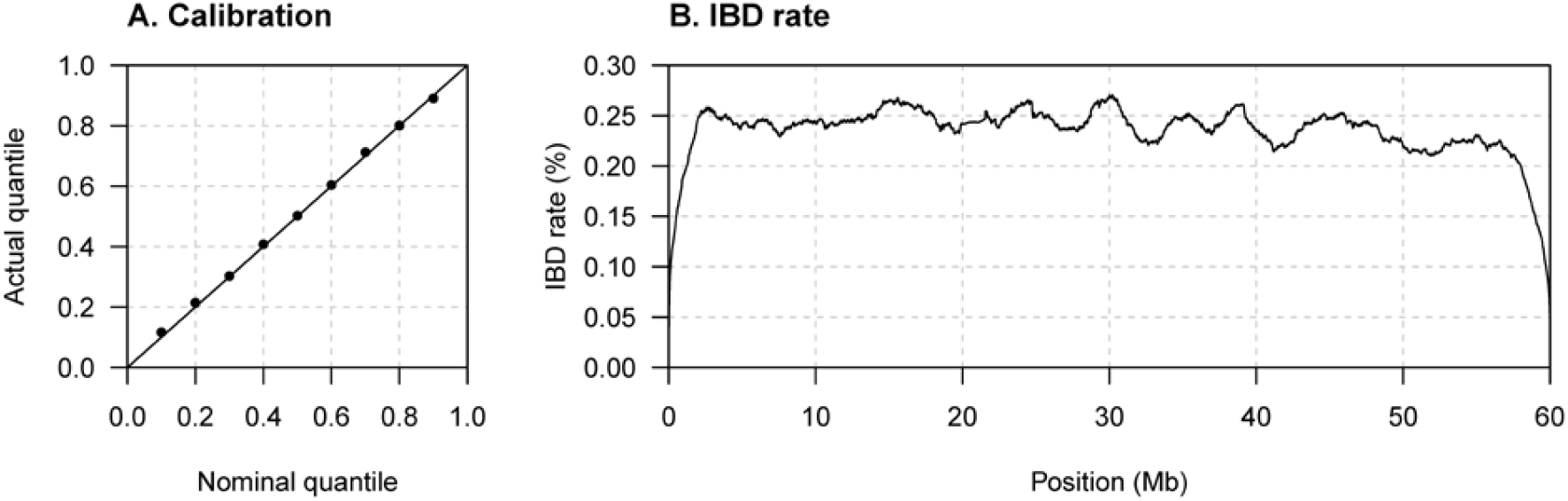
Method performance with uneven marker density. Sequence data on 2000 individuals were simulated under a constant effective population size. Markers located between 20 and 21.5 Mb were removed, and marker density varies every 100 kb (see Methods). The true haplotype phase is used in the analysis. (A) Quantile-quantile plot assessing the calibration of the estimated endpoint uncertainty. The actual quantile (y-axis) corresponding to a given nominal quantile (x-axis) is the proportion of segments for which the reported nominal quantile of the right endpoint is greater than the true right endpoint (points on the plot). The *y* = *x* line is shown for comparison. Results for the left endpoints are similar but are not shown. (B) The y-axis is the IBD rate, which is the percentage of pairs of haplotypes for which the position on the chromosome is covered by a sampled IBD segment with length > 2 cM for the haplotype pair. The IBD rate is calculated at 10 kb intervals.

Our method currently does not give special treatment to chromosome ends. A significant number of IBD segments terminate at the chromosome end, but our method does not assign any positive probability to an IBD segment ending exactly at the end of the chromosome. For analyzed segments with true right endpoint at the right end of the chromosome in these data, 25% have sampled right endpoint within 2 kb of the chromosome end, and 67% have sampled right endpoint within 100 kb of the chromosome end, but none have sampled right endpoint exactly at the chromosome end.

The ibd-ends analyses of these data used the default error rate of 0.0005. The error rate estimated by ibd-ends was 0.00039.

### UK-like simulated data

Figure 3 shows results for the simulated UK-like data with a 0.02% genotype error rate. The estimated uncertainty is well-calibrated (upper row of Figure 3), even when using inferred haplotype phase and when using data thinned to represent a 500k SNP array. As expected, endpoint uncertainty, as measured by the difference between the endpoint sampled from the uncertainty distribution and the true endpoint, is much higher when analyzing SNP array data rather than full sequence data (lower row of Figure 3, right vs left and middle columns). Results are also accurate with smaller sample sizes (200 or 1000 individuals’ Figure S1). When we removed markers to form a 1.5 cM gap (at 20-21.5 Mb), we found no noticeable change in calibration, and no excess rate of IBD in the region of the gap (Figure S2). We performed further analyses to evaluate endpoint estimation accuracy with a higher genotype error rate (Figure S3) and with different MAF thresholds (Figure S4). The results are well-calibrated with the higher genotype error rate (0.1%) when using true haplotypes. When using inferred haplotypes, the higher genotype error rate results in phase errors that reduce accuracy. The results are not particularly sensitive to the choice of MAF, but some miscalibration is observed when very rare variants are included (0.01% MAF).

**Figure 3:**
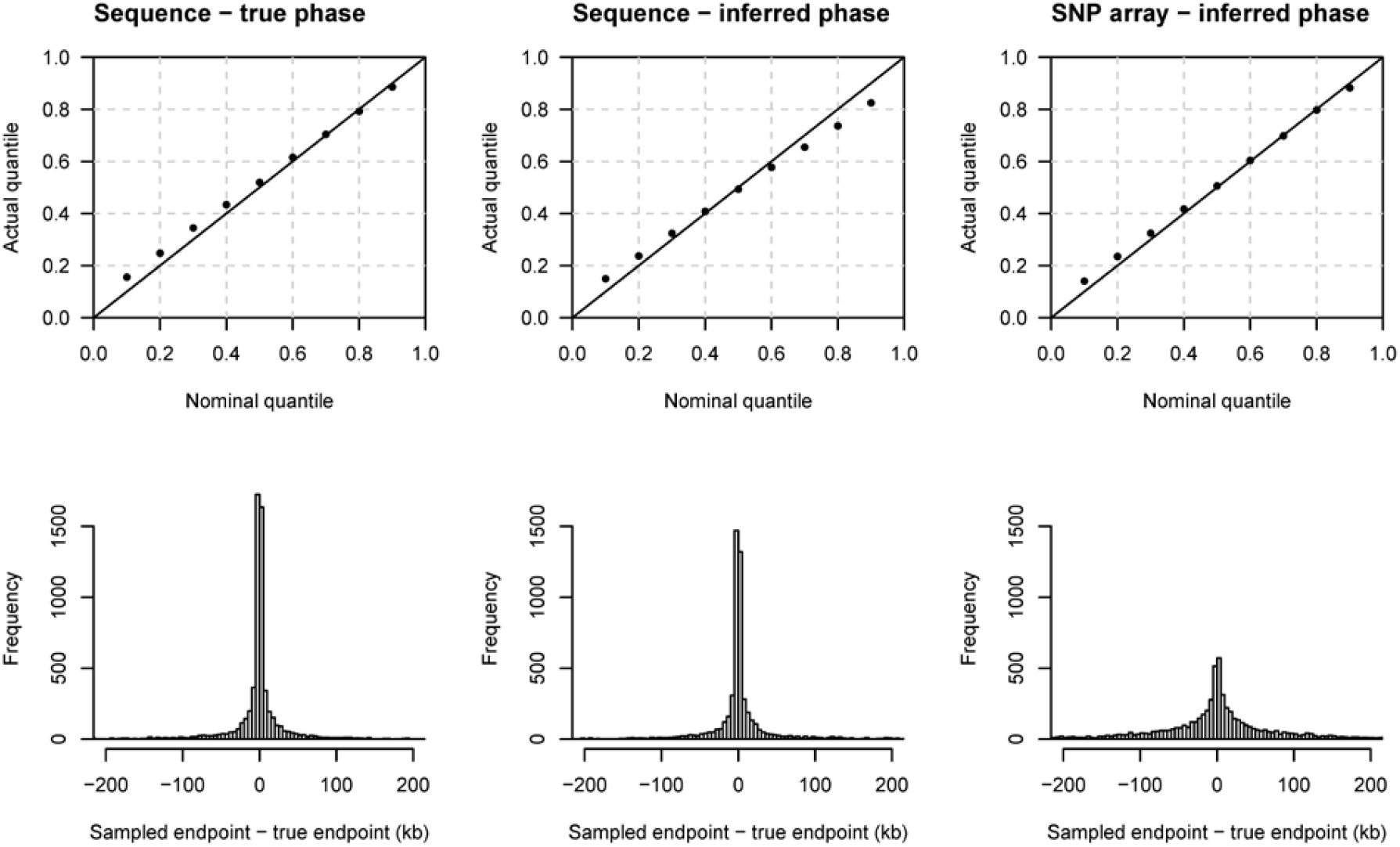
Method performance on UK-like simulated sequence and SNP array data. The data comprise 50,000 individuals simulated from a UK-like demographic history (see Methods), with a genotype error rate of 0.02%. True IBD segment endpoints were determined for 1000 individuals, and these individuals were used to generate the results in this figure. The top row shows quantile-quantile plots which assess the calibration of the estimated endpoint uncertainty. The *y* = *x* line is shown for comparison. The actual quantile (y-axis) corresponding to a given nominal quantile (x-axis) is the proportion of segments for which the reported nominal quantile of the right endpoint is greater than the true right endpoint. The bottom row shows histograms of the right endpoint sampled from the estimated posterior distribution minus the true right endpoint of the underlying segment. Histogram bin widths are 5 kb. Results for the left endpoints are similar but are not shown. The left column is for analysis using the true haplotype phase. The middle column is for analysis using haplotype phase inferred using Beagle 5.1. The right column is for data thinned to match a SNP array with 500,000 markers genome-wide (10,000 markers in the simulated 60 Mb interval), and with haplotype phase inferred using Beagle 5.1.

We found that using an analyses error rate that is up to three times higher or lower than the estimated error rate gave accurate results (Figure S5 and Table S1). When using an analysis error rate that is outside this range, the estimated error rates produced by ibd-ends are within the range of error rates that will provide good results in a subsequent analysis (Table S1). If the estimated error rate differs from the analysis error rate by more than three-fold, we recommend redoing the analysis using the estimated rate. Pilot results on a small chromosome can be used to determine whether the analysis error rate needs to be changed from the default value.

Compute times for ibd-ends analyses with default settings were 1.4 hours for the full UK-like data with 50,000 individuals (17 million IBD segments), and 0.5 hours for 1000 individuals (7000 IBD segments) using a 24-core compute node with 24 Intel Xeon Silver 4214 2.2 GHz processors and 382 GB of memory.

### Heterogeneous simulated data

In the heterogeneous simulation, half of the simulated individuals are from a population with an African demographic history, while the other half are from a population with a European demographic history (see Methods). Analyzing these data together violates the assumptions of the endpoints modelling, however the results are not excessively mis-calibrated (Figure S6A). For example, 18% of the true endpoints are closer to the center of the IBD segment than the nominal 10^th^ percentile. When analyzing the African individuals separately (Figure S6B) or the European individuals separately (Figure S6C), the results are well-calibrated.

### UK Biobank data

We sampled one left endpoint and one right endpoint from the endpoint uncertainty distributions of 77.7 billion candidate autosomal IBD segments of length 2 cM or larger that hap-IBD detected in the 408,883 White British UK Biobank individuals. We refer to the segment defined by the sampled end points as the “sampled IBD segment”. Analysis was parallelized by chromosome. Total wall clock computing time across all chromosomes was 7.5 hours for hap-ibd and 150 hours for ibd-end using a 24-core Intel Xeon Silver 4214 2.2 GHz compute node. The estimated error rates from each chromosome varied from 0.00027 to 0.00034 when using the default analysis error rate of 0.0005.

There were 51.7 billion sampled IBD segments with length > 2 cM. Every 10 kb along each chromosome we computed the number of sampled IBD segments with length > 2 cM covering the position (Figure 4). The IBD rate is the number of IBD segments covering a position divided by the number of haplotype pairs. Each individual contributes two haplotypes, and all haplotype pairs are considered except those pairs within the same individual. A high rate of IBD at a position is a signal of possible recent strong natural selection.^14–16^

**Figure 4:**
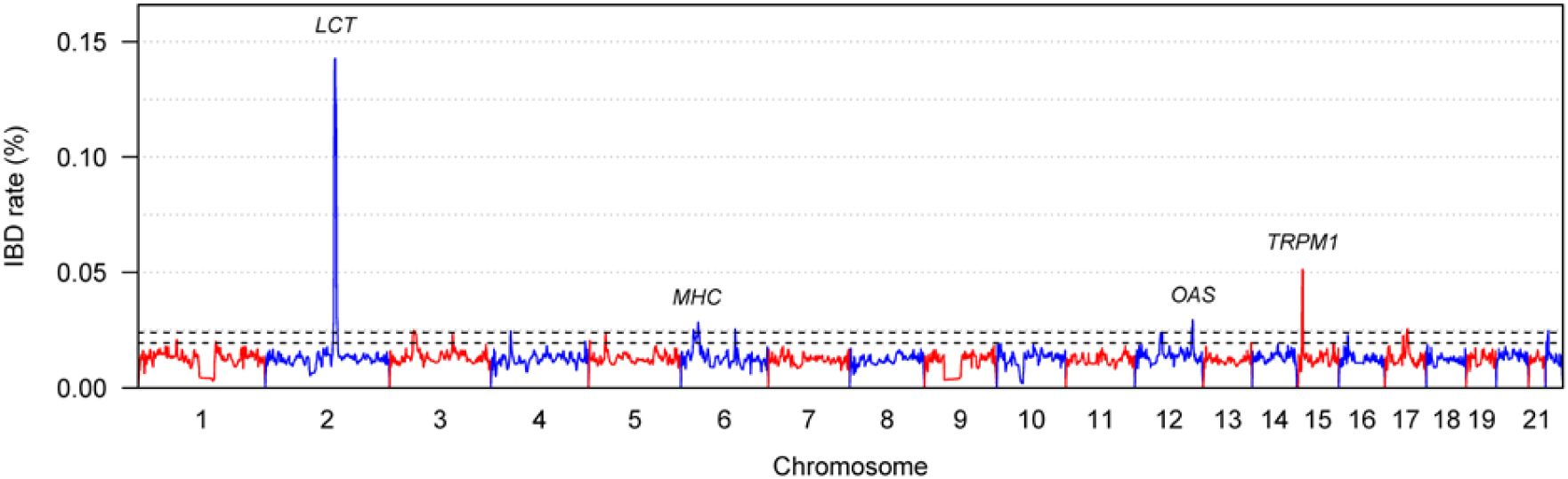
Rate of IBD segments along the autosomes in UK Biobank White British data. The x-axis shows position along each chromosome. Chromosomes alternate in color. Notable genes and regions (*LCT, MHC, OAS*, and *TRPM1*) located within the four highest peak regions are labelled. The y-axis is the IBD rate, which is the percentage of pairs of haplotypes for which the position on the chromosome is covered by a sampled IBD segment with length > 2 cM for the haplotype pair. The IBD rate is calculated at 10 kb intervals. The black dashed lines show the thresholds of 0.024% and 0.0196% used for the results in Tables 1 and 2 respectively.

The median IBD rate is 0.0126%. There are ten regions with an IBD rate higher than 0.024% (Table 1). Nine of these are genomic regions known to be undergoing significant levels of selection, indicating the success of this approach in finding real signals of selection. Four of the regions have been shown to have adaptive introgression from Neanderthals (*OAS, CCR9/CXCR6, TLR1/6/10*, Type II Keratins). Five of the regions of selection play a role in immunity (*MHC, OAS, CCR9/CXCR6, TLR1/6/10, PRDM1*). Other regions are involved in nutrition (*LCT*), skin and hair traits (Type II Keratins and *TRMP1*), and fertility (*MAPT* inversion).

**Table 1:** Regions of highest IBD rate in UK Biobank White British analysis. Regions in which the maximum IBD percentage is at least 0.024% are shown. Positions are in GRCh37 coordinates.

| Chromosome | Peak position (Mb) | Region (Mb)<sup>a</sup> | Max IBD % | Notes<sup>b</sup> |
| --- | --- | --- | --- | --- |
| 2 | 136.65 | 134.75-138.47<br>(2q21-22) | 0.1426 | *LCT* (136.55-136.59) is known to be subject to recent positive selection.<sup>50</sup> |
| 3 | 47.62 | 45.18-53.22<br>(3p21) | 0.0249 | Chemokine receptor genes including *CCR9* (45.93-45.94) and *CXCR6* (45.98-45.99). This region shows adaptive introgression from Neanderthals.<sup>65</sup> |
| 4 | 38.84 | 38.12-40.21<br>(4p14) | 0.0246 | Toll-like receptor genes including *TLR1*, *TLR6*, and *TLR 10* (38.77-38.86). This locus is known to be under positive selection.<sup>69</sup> |
| 6 | 24.87<br>33.97 | 24.03-36.63<br>32.43-36.27<br>(6p21-22) | 0.0252<br>0.0284 | MHC locus containing *HLA* genes (28.48-33.45) is known to be subject to strong pressures of natural selection.<sup>54</sup> |
| 6 | 106.33 | 105.98-107.31<br>(6q21) | 0.0255 | *PRDM1* (106.53-106.56). This locus shows a signal of recent selection.<sup>64</sup> |
| 12 | 53.46 | 49.17-54.75<br>(12q13) | 0.0240 | Type II Keratins (*KRT1-8*; 52.63-53.32). This locus has experienced adaptive introgression from Neanderthals.<sup>66</sup> |
| 12 | 113.44 | 111.64-114.11<br>(12q24) | 0.0294 | *OAS* locus (*OAS1-3*) (113.34-113.45). This locus has experienced adaptive introgression from Neanderthals.<sup>59</sup> |
| 15 | 31.61 | 30.90-32.46<br>(15q13) | 0.0513 | *TRPM1* (31.29-31.45). A pigmentation gene under selection in non-Africans.<sup>56</sup> |
| 17 | 44.78 | 42.01-45.89<br>(17q21) | 0.0256 | 17q21.31 *MAPT* inversion (43.44-44.85).<sup>c</sup> H2 form has been positively selected in Europe.<sup>60</sup> |
| 22 | 22.56 | 21.36-23.05<br>(22q11) | 0.0248 | *UBE2L3* (21.90-21.98) is associated with multiple auto-immune diseases.<sup>72</sup> |

<sup>a</sup> Region in which the IBD percentage is at least 80% of the value at the peak.
<sup>b</sup> See the main text for further notes and references.
<sup>c</sup> Positions from Zody et al.<sup>79</sup> lifted over from build 36 (chromosome 17: 40.80-42.20 Mb) to build 37.

The highest selection signal, with an IBD rate of 0.14%, comes from a chromosome 2 region containing the *LCT* gene which has a variant selected for lactose tolerance in Europeans.^50^ The selected variant is thought to have arisen, or at least begun to increase in frequency, around the time of the advent of cattle farming in Europe, around 7500 years ago,^51^ and selection has been so strong that the selected variant allele frequency is now around 75% in individuals of British descent.^52^ Since IBD is affected by recent selection, it is not surprising that this signal is most prominent.

In contrast, the immunity-related *HLA* genes in the multi histocompatibility complex (*MHC*) on chromosome 6 have been under selection over a much longer time period,^53; 54^ and this region has a much lower peak IBD rate (0.028%) than the *LCT* region. Various sub-regions within the *MHC* appear to have been subject to adaptive introgression from Neanderthals and Denisovans, but it is difficult to be certain because long-term balancing selection across the region can result in signals that look like adaptive introgression.^55^

The second-highest signal, with an IBD rate of 0.051%, comes from a chromosome 15q13 region containing *TRPM1*, a pigmentation gene that has been shown to have been subject to selection in non-Africans.^56; 57^

The high IBD rate region on chromosome 12q24 (0.029% IBD rate) encompasses the *OAS* locus which is involved in immunity.^58^ This locus has a Neanderthal haplotype present at high frequency in non-Africans that has been subject to positive selection.^59^

The high IBD rate region on chromosome 17q21 (0.026% IBD rate) encompasses the 17q21.31 *MAPT* inversion, for which the H2 form has undergone positive selection in Europe and is associated with increased fertility and higher recombination rates in females in Iceland.^60^

The high IBD rate region on chromosome 6q21 (0.026% IBD rate) contains the *PRDM1* and *ATG5* genes. This pair of genes is associated with autoimmune diseases and cancer^61–63^ and shows a signal of recent selection in HapMap data.^64^

The high IBD rate region on chromosome 3p21 (0.025% IBD rate) is a region of adaptive introgression from Neanderthals.^65; 66^ It contains chemokine receptor genes, including CCR9 and CXCR6, that are involved in immunity.^67; 68^

The high IBD rate region on chromosome 4p14 (0.025% IBD rate) contains several toll-like receptor genes (*TLR1, TLR6*, and *TLR10*) that are involved in immunity, and this region has experienced adaptive introgression from Neanderthals.^66^ As well as the adaptive introgression, this region shows other signals of selection, including geographic differentiation within the UK^69^ and signs of recent positive selection among non-Africans.^70^

The high-IBD rate region on chromosome 12q13 (0.024% IBD rate) contains the Type II Keratin genes, which code filament proteins that provide a major structural role in epithelial cells.^71^ This region has experienced adaptive introgression from Neanderthals.^66^

The remaining locus on chromosome 22q11 (0.025% IBD rate) has not previously been highlighted as being under selection, to the best of our knowledge. This locus includes *UBE2L3* which is associated with multiple auto-immune diseases.^72^ These associations make this locus a strong candidate for natural selection.^73^

We also investigated the next ten regions with highest IBD rates (Table 2). These regions include the pigmentation gene *SLC45A2* on chromosome 5p13 (0.024% IBD rate) and the epidermal differentiation complex locus on chromosome 1q21.3 (0.020% IBD rate). Both of these loci are known to have undergone recent positive selection.^74; 75^

**Table 2:** Secondary regions from UK Biobank selection scan. Regions with maximum IBD rate between 0.0195% and 0.024% are shown. Positions (in Mb) are in GRCh37 coordinates.

| Chromosome | Peak position (Mb) | Region (Mb)<sup>a</sup> | Max IBD % | Notes |
| --- | --- | --- | --- | --- |
| 1 | 76.49 | 74.93-77.33<br>(1p31.1) | 0.0209 | *ST6GALNAC3* (76.54-77.10) is subject to virus-driven selective pressure.<sup>80</sup> |
| 1 | 152.78 | 151.58-153.75<br>(1q21.3) | 0.0201 | Epidermal differentiation complex locus (151.96-153.60).<sup>81</sup> The locus with the greatest differentiation between humans and chimpanzees.<sup>82</sup> Recent positive selection in humans.<sup>75</sup> |
| 3 | 123.41 | 122.36-124.70<br>(3q21) | 0.0233 | *ADCY5* (123.00-123.17) is associated with birth weight<sup>83</sup> and type II diabetes,<sup>84</sup> and is under long term balancing selection.<sup>85</sup> |
| 4 | 184.59 | 184.30-185.48<br>(4q35.1) | 0.0201 |  |
| 5 | 2.68 | 2.32-2.99<br>(5p15.33) | 0.0205 |  |
| 5 | 33.84 | 32.56-34.63<br>(5p13) | 0.0239 | *SLC45A2* (33.94-33.98, previously known as *MATP*) is a pigmentation gene that has undergone recent positive selection in Europe.<sup>74;</sup><br><sup>86</sup> |
| 13 | 112.87 | 112.54-113.55<br>(13q34) | 0.0196 | *SPACA7* (113.03-113.09) is a reproduction-related gene under positive selection in primates.<sup>87</sup> |
| 16 | 18.08 | 17.35-18.74<br>(16p12.3) | 0.0229 |  |
| 17 | 36.03 | 35.17-36.51<br>(17q12) | 0.0226 | *HNF1B* (36.05-36.10, previously known as *TCF2*) is associated with diabetes.<sup>88; 89</sup> |
| 22 | 18.89 | 18.57-19.60<br>(22q11.21) | 0.0199 |  |

<sup>a</sup> Region in which the IBD percentage is at least 80% of the value at the peak.

The variability of IBD rate across the genome is much lower for ibd-ends than for hap-ibd (Figure S7). The standard deviation of the ibd-ends IBD rate is 0.0056%, whereas the standard deviation for hap-ibd is an order of magnitude higher at 0.060%. Ibd-ends is a much better tool for investigating regions of potential selection than length-based methods because length-based methods can report artifactually high rates of IBD in some regions, such as regions containing large gaps in marker coverage. For example, IBD segment detection with four length-based IBD detection methods produce a 40-3000 fold higher IBD rate at the chromosome 1 centromere in UK Biobank data compared to the background rate.^33^

Misspecification of the genetic map can also lead to spurious regions of high IBD rate in IBD selection scans, because overestimation of genetic length will result in a larger number of IBD segments that pass the length threshold. To investigate the impact of map choice on the IBD selection analysis, we analyzed a subset of 50,000 White British individuals with three maps. We observed considerable differences in the regions of highest IBD rate when using different maps (Figure S8). With the HapMap linkage-disequilibrium (LD) based map^76^ there are 20 regions with IBD rate > 0.05% (compared to two with Bherer et al.’s family-based map),^49^ with three of these occurring at the locations of centromeres. Furthermore, the IBD rate has higher variability when using the HapMap map, with a standard deviation of 0.018% (compared to 0.0057% with Bherer et al.’s map in this 50,000-individual subset). LD-based maps are known to be biased in regions of selection, with genetic distances underestimated in these regions,^77^ which would lead to an apparent decrease in IBD rate in these regions. Thus, increases in apparent IBD rate with the LD-based HapMap map compared to Bherer et al.’s pedigree-based map must be due to other effects, such as unmodelled features of the data. The IBDrecomb map is designed to obtain uniform IBD rate across the genome.^13^ Thus regions of likely selection such as *LCT* do not have high IBD rates when using this map. The analysis with IBDrecomb still has some regions of high IBD rate (19 regions with IBD rate > 0.05%), which are concentrated at chromosome ends (where inflation of map distances is known to occur^13^) and at centromeric gaps. These regions are likely to be artifacts.

The MHC region has a much higher selection signal from analysis with the HapMap map than with Bherer et al.’s map. The genetic lengths of the MHC are 3.29 cM for the HapMap map, 2.20 cM for Bherer et al.’s map and 1.44 cM for the IBDrecomb map. Three previous IBD-based selection analyses have used the HapMap map,^14–16^ and two of these analyses also had very strong MHC signals.^14; 16^

## Discussion

We presented a method for calculating the posterior probability distributions of IBD segment endpoints. We showed that the method can be applied to large data sets, such as the UK Biobank SNP array data on 408,883 White British individuals and simulated sequence data on 50,000 individuals. In the UK Biobank data, we analyzed 77.7 billion candidate IBD segments, and found 51.7 billion IBD segments for which the length based on sampled endpoints is greater than 2 cM.

In addition to quantifying endpoint uncertainty, a major advantage of our method is that it handles genotype errors and other discordances within IBD segments in a principled way, in contrast to many other methods for IBD segment detection which use ad hoc approaches. Our method does not directly account for haplotype phase uncertainty, however statistical phasing of non-rare variants in large array-typed cohorts is now extremely accurate,^36^ and technologies for generating highly-accurate, phase-resolved sequence data are becoming available.^37^

The sampled lengths and endpoints of the segments are unbiased and have low variance. IBD segments defined by the sampled endpoints can be used in downstream analyses. In addition, the estimates of endpoint uncertainty can be used to include more data in analyses. For example, when estimating genome-wide mutation rates from IBD segments, it is important to be confident that one does not count mutations that actually lie outside the IBD segment. In the past, this has been achieved by trimming 0.5 cM from the putative IBD endpoints,^10; 12^ but trimming using a small quantile of the uncertainty distribution would be expected to result in less trim (and hence more data), while maintaining accuracy. Alternatively, methods could be developed that directly account for endpoint uncertainty without trimming.

Another analysis that could incorporate estimated IBD segment end point uncertainty is IBD-based estimation of recombination maps. Endpoints of IBD segments are points of past recombination which are used to estimate the map. Consequently, misspecification of IBD endpoints adds noise and bias to the resulting map. One could improve recombination map estimation by incorporating endpoint uncertainty into an iterative procedure that uses the current estimated recombination map to refine the estimates of IBD segment endpoints.

IBD segment endpoint uncertainty could also be used to improve IBD-based estimation of recent demographic history. For this application, it is not the actual IBD endpoints that matter, but rather the distribution of IBD segment lengths, which one could estimate by sampling endpoints from the posterior distribution. A threshold of 2 or 3 cM on IBD segment length is usually applied with current methods since lengths of shorter segments have higher relative uncertainty.^4–8^ With sampled endpoints, it may be possible to use a lower length threshold and thus estimate demographic history further into the past.

We found that the IBD segments obtained from sampled endpoints provide excellent input data for an IBD-based selection scan. Nine of the ten top regions in our UK Biobank analysis are regions of known selection, and the remaining region is a good candidate for selection. Two particular features of our IBD-based selection analysis contribute to its success. The first is unbiased estimation of IBD segment lengths, even in the presence of gaps in marker coverage, which eliminates many spurious signals. In contrast, length-based IBD detection methods tend to have regions with inflated IBD rates,^33^ which would cause spurious signals if used in a selection scan. The second is the choice of genetic map. We found that for IBD-based selection scans, pedigree-based maps based on actual observations of recombination produce more accurate results than maps based on LD or on IBD sharing, because effects of selection and other unmodelled features in local genomic regions can bias LD-based and IBD-based maps, and this bias results in spurious signals of selection.

Two limitations of the current study could be addressed in future work. First, our modelling assumes a homogeneous population with the same distribution of IBS segment lengths for all pairs of individuals.

Our analysis of a simulated combined African and European ancestry sample indicates that violation of this assumption introduces a little mis-calibration in the posterior endpoint probabilities. One solution would be to adjust for ancestry of the samples, or for local ancestry in the case of admixed samples. Second, we did not give any special treatment to the ends of chromosomes. Consequently, IBD segments that truly extend to the end of the chromosome are estimated to have endpoints occurring near, rather than at, the chromosome ends. One work-around would be to use the input candidate segment endpoint in preference to the ibd-ends endpoint when the input candidate segment endpoint is located at the chromosome end. A more sophisticated solution would be to use the estimated distribution of IBD segment lengths as a prior distribution when analyzing IBD segments that may extend to the end of the chromosome.

## Appendices

## Appendix 1: Prior for IBD length distribution

In this appendix we derive a formula for the prior probabilities *P*(*R* ∈ (*x_i_*, *x*_*i*+1_)) in Equation 1. Let *Y* = *R* – *L*_0_ be the length (measured in Morgans) from the left endpoint, *L*_0_, of the segment to the right endpoint *R*. Here we assume that the left endpoint is known, but in practice we iteratively update its estimated value as described in the main text. We write *F*(*y*) = *P*(*Y* ≤ *y*) for the prior probability distribution on *Y*. We model the population size as constant diploid size *N*, with *N* = 10,000, which reflects the approximate average historical size of out-of-Africa populations.

Summing over possible values for *G*, the number of generations to the common ancestor of the IBD segment, we obtain

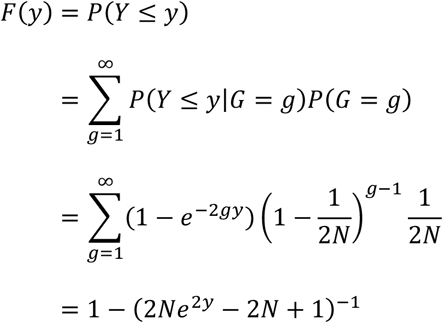

In these calculations, we use the fact that the IBD segment length conditional on the number of generations *g* to the common ancestor is exponentially distributed with rate parameter 2*g*,^7^ and the fact that the number of generations to the most recent common ancestor in a population of constant size *N* is geometrically distributed.^4^ The final expression for *F*(*y*) in the calculation is obtained by applying the formula for a geometric series.

The prior distribution for the right endpoint is:

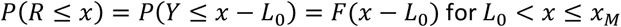

and the probability in Equation 1 that *R* falls in a certain interval can be calculated as:

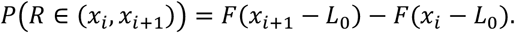

In Appendix 3, we will require the inverse of *F*, which is 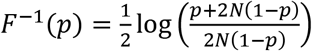.

## Appendix 2: Modelling the IBS data beyond the IBD segment

In this appendix we describe how to calculate the conditional probability *P*(*D*[*i* + 1, *M*] |*R* ∈ (*x_i_*, *x*_*i*+1_)) in Equation 1. This is the probability of the IBS data to the right of the right IBD endpoint.

Let *m_i_*(*J*) (*j* ≥ 1) be the ordered indices of the discordant markers to the right of *x_i_*, plus the last marker on the chromosome if that marker is not included in the list of discordant positions. We approximate the IBS process to the right of the IBD endpoint as a renewal process with a renewal every time there is a discordant marker. Then the probability of the IBS data to the right of *x_i_* given that the IBD endpoint occurred in the interval (*x_i_*, *x*_*i*+1_) is:

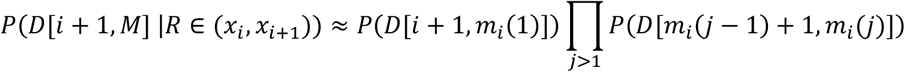

Each interval of IBS data in the preceding equation has the form *D*[*a, b*], where *a* and *b* are marker indices, and has the property that the alleles on the two haplotypes *H*_1_ and *H*_2_ are identical at all positions *i* satisfying *a* ≤ *i* < *b*, but are discordant at position *b* (except in some cases where *b* is the last marker on the chromosome). Our approach to estimating the probabilities of these IBS interval data is to estimate the probability *p_a,b_* that two randomly chosen haplotypes have identical alleles over the interval [*a, b*] (i.e. for all marker indicies *i* with *a* ≤ *i* ≤ *b*). If haplotypes *H*_1_ and *H*_2_ have identical alleles at marker *b*, then we set *P*(*D*[*a, b*]) = *p_a,b_*, and if the two haplotypes have discordant alleles at marker *b*, we set *P*(*D*[*a, b*]) = *p_a,(b−1)_ – p_a,b_*. We estimate *p_a,b_* empirically from the data.

Our method for estimating *p_a,b_* depends on the length of the interval [*a, b*]. For short intervals, we use data in the interval so that the estimated probability incorporates the local genomic context, such as high or low marker density, high or low heterozygosity, and high or low levels of LD. For long intervals, we will use chromosome-wide data. As the interval length becomes longer, *p_a,b_* will tend to decrease because the probability of observing a long IBS interval for a random pair of haplotypes is small. Small probabilities are more difficult to estimate than long ones, so we need to bring in data from the rest of the genome. Furthermore, long intervals represent long IBS (alleles at all markers in the interval are identical except for the final marker), and long IBS is likely to be the result of a long IBD segment. Excluding the effects of selection, the distribution of IBD lengths should be uniform across the genome, so estimating the frequency of such long IBS segments from data across the genome is appropriate.

For short intervals, we estimate *p_a,b_* by the proportion of pairs of haplotypes that have identical by state alleles for markers in the interval [*a, b*]. We consider an interval [*a, b*] to be a short interval if the estimated 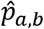 satisfies 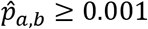 when it is estimated from the haplotypes in the interval. We use 10,000 randomly sampled haplotypes (or all haplotypes if the sample size is 5000 or fewer individuals) to estimate the short interval IBS probabilities.

For longer intervals we estimate *p_a,b_* using the global distribution of sampled one-sided IBS lengths. A one-sided IBS length is the cM distance from a random starting position to the first non-IBS marker in the direction toward the center of the chromosome for a random pair of haplotypes. We estimate *p_a,b_* by the proportion of sampled one-sided IBS lengths that are greater than (*x_b_* – *x_a_*). When estimating the global IBS length, we randomly select 1000 positions and randomly sample 2000 one-sided IBS lengths for each position. We then exclude the sampled lengths at positions for which the IBS length are significantly longer than average. We do this by calculating the 90th percentile of the 2000 segment lengths for each position. If the 90^th^ percentile at a position is more than 3 times the median 90th percentile from all 1000 positions, we discard the position. This filtering protects against selecting positions near gaps in marker coverage.

## Appendix 3: Obtaining the posterior cumulative distribution function for the endpoint

In this appendix we describe how to calculate the cumulative distribution function of the posterior distribution for the right endpoint of the IBD segment, as well as how to sample from this posterior distribution.

We assume that the data *D* are not informative about the location of the endpoint within the inter-marker interval given that the endpoint occurs within the interval, i.e. we assume that *P*(*x_i_* < *R* ≤ *x*|*x_i_* < *R* < *x*_*i*+1_, *D*) = *P*(*x_i_* < *R* ≤ *x*|*x_i_* < *R* < *x*_*i*+1_) for *x_i_* < *x* < *x*_*i*+1_, since there are no data within the interval. Write *p_i_* = *P*(*R* ≤ *x_i_*|*D*), which can be estimated using Equation 1 with the procedures described above. As in Appendix 1, we write *Y* = *R* – *L*_0_ for the length of the segment (in Morgans), and *F*(*y*) = *P*(*Y* ≤ *y*) for the prior on IBD lengths. For *x* satisfying *x_i_* < *x* < *x*_*i*+1_:

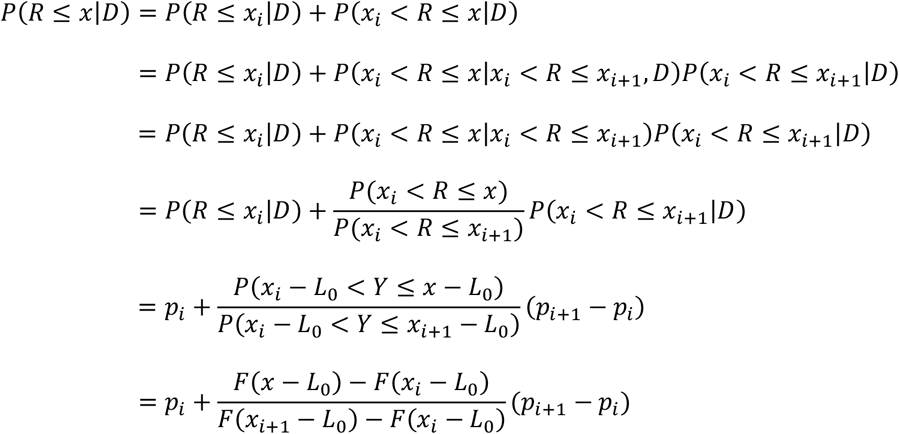

Note that *p_i_* = *P*(*R* ≤ *x_i_*|*D*) is conditional on the data, whereas *F*(*x_i_* – *L*_0_) = *P*(*Y* ≤ *x_i_* – *L*_0_) = *P*(*R* ≤ *x_i_*) is a prior probability and is not conditional on the data.

Then to find the *p*-th quantile we want to find *x*^(*p*)^ such that *P*(*R* < *x*^(*p*)^|*D*) = *p*. Solving the above equation for *x* = *x*^(*p*)^ we obtain

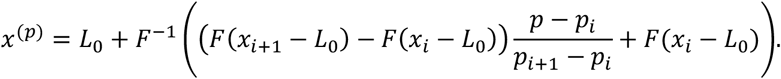

The formula for *F*^−1^(*p*) can be found in Appendix 1.

In order to obtain a sampled value from the posterior probability of the endpoint, we first generate a realization *u* from the Uniform(0,1) distribution, and we then obtain *x*^(*u*)^ using the above formula with *u* in place of *p*, which is then a realization from the desired distribution. This is an example of using the inverse transform principle to sample from a distribution.^78^

## Supplemental Data

Supplemental data consist of 8 figures and 1 table.

## Acknowledgments

Research reported in this publication was supported by the National Human Genome Research Institute of the National Institutes of Health under award numbers HG005701 and HG008359. The content is solely the responsibility of the authors and does not necessarily represent the official views of the National Institutes of Health. This research has been conducted using the UK Biobank Resource under Application Number 19934.

## Web Resources

ibd-ends: https://github.com/browning-lab/ibd-ends

Bherer et al.’s refined European map: https://github.com/cbherer/Bherer_etal_SexualDimorphismRecombination/blob/master/Refined_EUR_genetic_map_b37.tar.gz

HapMap map: ftp://ftp.ncbi.nlm.nih.gov/hapmap/recombination/2011-01_phaseII_B37/genetic_map_HapMapII_GRCh37.tar.gz

IBDrecomb European-American map: https://github.com/YingZhou001/IBDrecomb/tree/master/maps.b37/FHS.map.tar.gz

---

**Figure S1:**
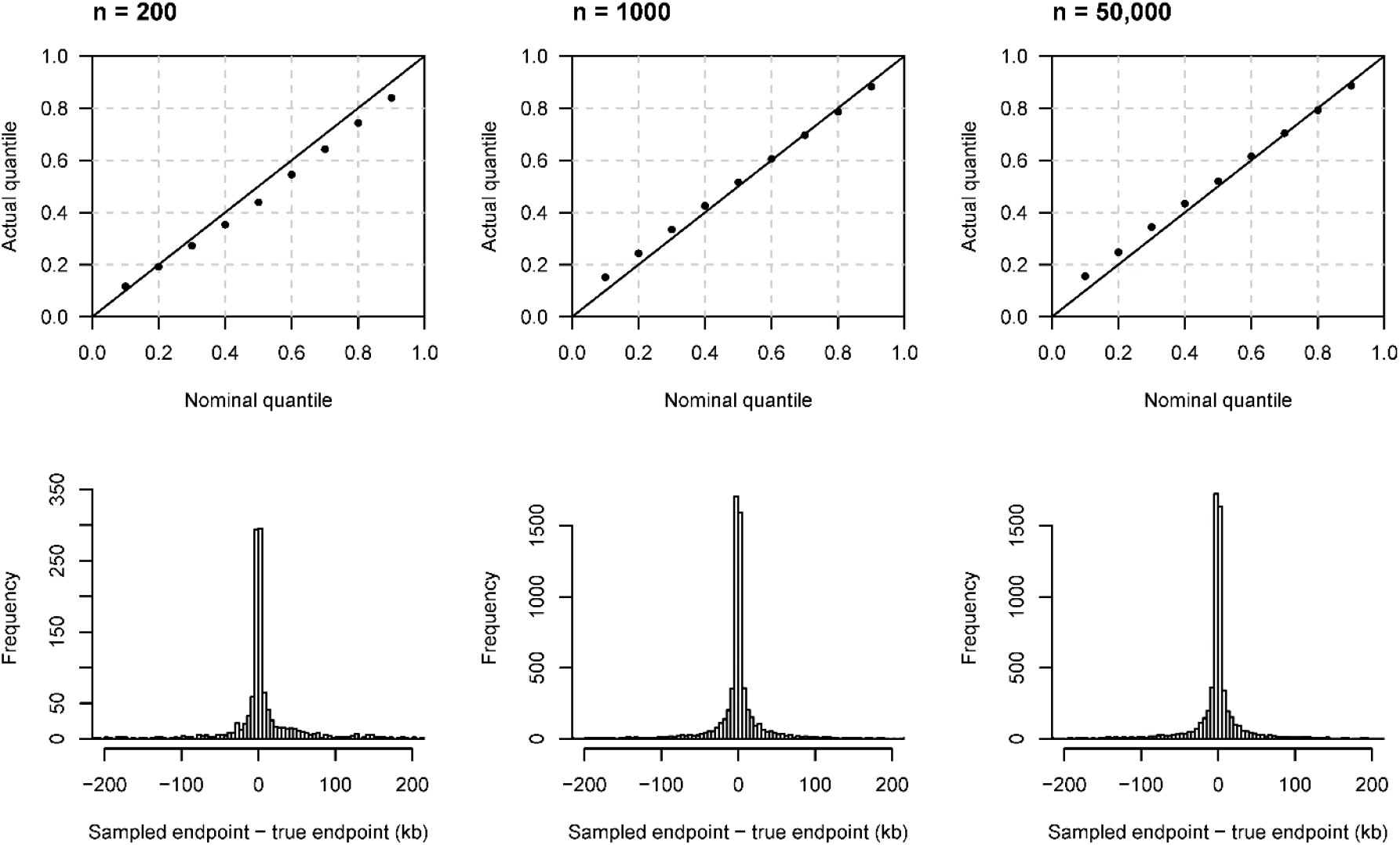
Performance on UK-like simulated data with varying sample sizes. The input data comprise sequence data on individuals simulated from a UK-like demographic history (see Methods), with a genotype error rate of 0.02%, and true haplotype phase. The top row shows quantile-quantile plots which assess the calibration of the estimated endpoint uncertainty. The *y* = *x* line is shown for comparison. The actual quantile (y-axis) corresponding to a given nominal quantile (x-axis) is the proportion of segments for which the reported quantile of the right endpoint is greater than the true right endpoint. The bottom row shows histograms of the right endpoint sampled from the estimated posterior distribution minus the true right endpoint of the underlying segment. Histogram bin widths are 5 kb. Results for the left endpoints are similar but are not shown. The left column is for five analyses each using 200 individuals. The middle column is for analysis using 1000 individuals. The right column is for analysis using 50,000 individuals with results were assessed on 1000 individuals for which true IBD endpoints were determined; these results are the same as the left column in Figure 3. The five n=200 analyses (left column) altogether involve approximately 20% as many haplotype pairs as the other two analyses, and hence the results are subject to more statistical variation. Since there are 20% fewer data points than the n=1000 and n=50,000 analyses, the y-axis of the n=200 histogram (lower left) has been scaled proportionally, in order to make it comparable to the other two analyses.

**Figure S2:**
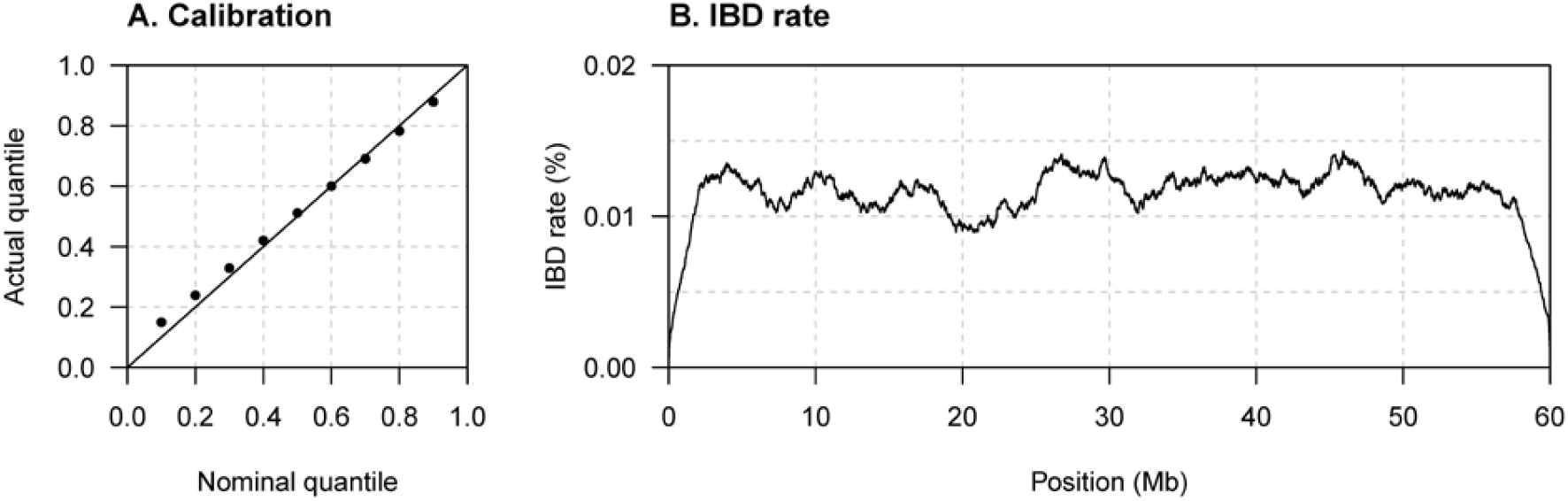
Method performance on simulated UK-like data with 1.5 Mb marker gap. 1000 individuals from the UK-like simulated sequence data with true phase were analyzed. Markers located between 20 and 21.5 Mb were removed. (A) Quantile-quantile plot assessing the calibration of the estimated endpoint uncertainty. The *y* = *x* line is shown for comparison. The actual quantile (y-axis) corresponding to a given nominal quantile (x-axis) is the proportion of segments for which the reported nominal quantile of the right endpoint is greater than the true right endpoint. Results for the left endpoints are similar but are not shown. (B) The y-axis is the IBD rate, which is the percentage of pairs of haplotypes for which the position on the chromosome is covered by a sampled IBD segment with length > 2 cM. The IBD rate is calculated at 10 kb intervals.

**Figure S3:**
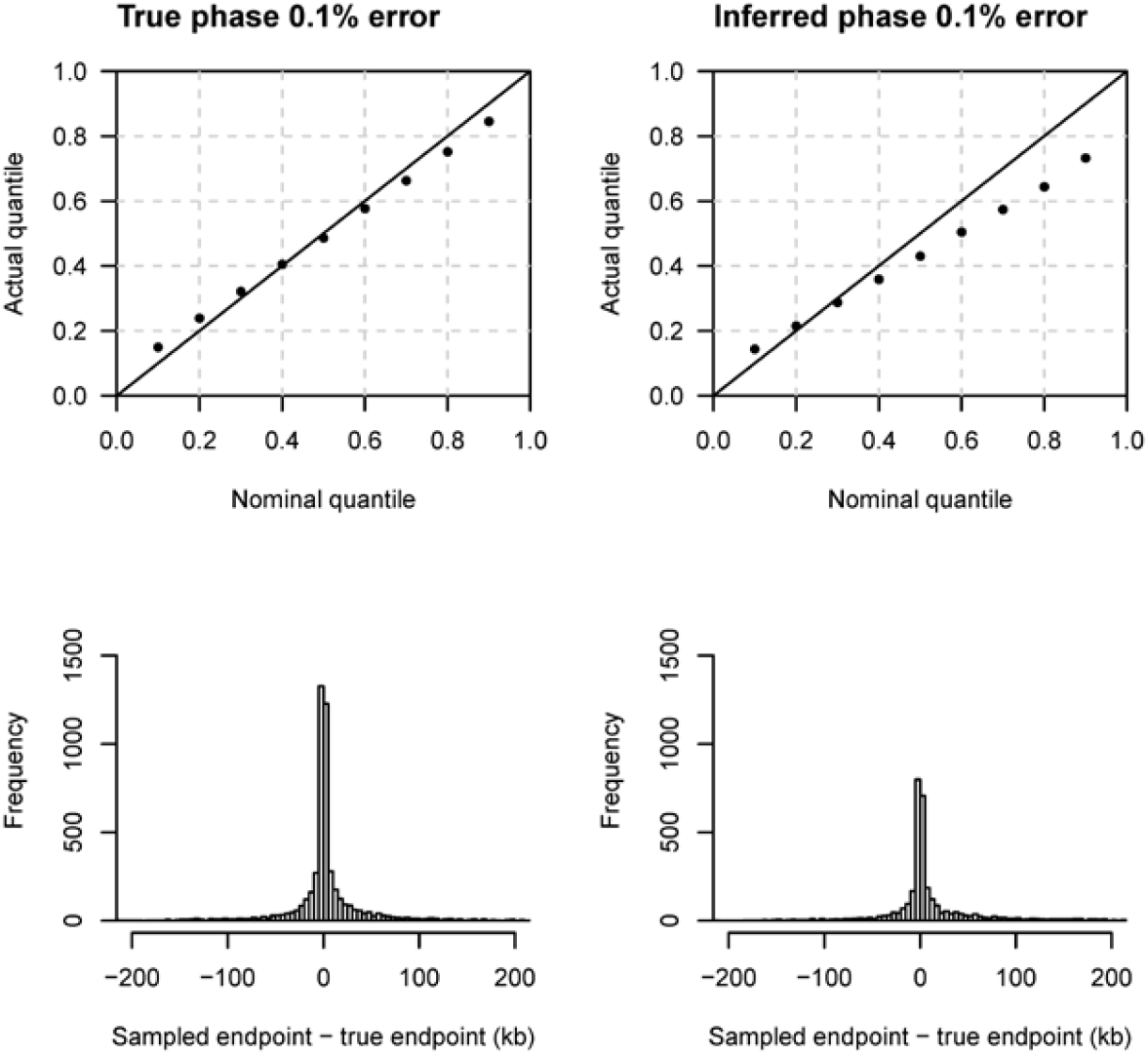
Performance on UK-like simulated data with a 0.1% rate of genotype error. The data comprise sequence data on 50,000 individuals simulated from a UK-like demographic history (see Methods), with a genotype error rate of 0.1%. Results were calculated using true IBD segment endpoints for all pairs of individuals within a subset of 1000 individuals. The top row shows quantile-quantile plots which assess the calibration of the estimated endpoint uncertainty. The *y* = *x* line is shown for comparison. The actual quantile (y-axis) corresponding to a given nominal quantile (x-axis) is the proportion of segments for which the reported nominal quantile of the right endpoint is greater than the true right endpoint. The bottom row shows histograms of the right endpoint sampled from the estimated posterior distribution minus the true right endpoint of the underlying segment. Histogram bin widths are 5 kb. Results for the left endpoints are similar but are not shown. The left column is for analysis using the true haplotype phase. The right column is for analysis using haplotype phase inferred using Beagle 5.1. There are 26% fewer segments in the inferred phase analysis, because the higher genotype error rate results in some undetected IBD segments at the hap-ibd detection step.

**Figure S4:**
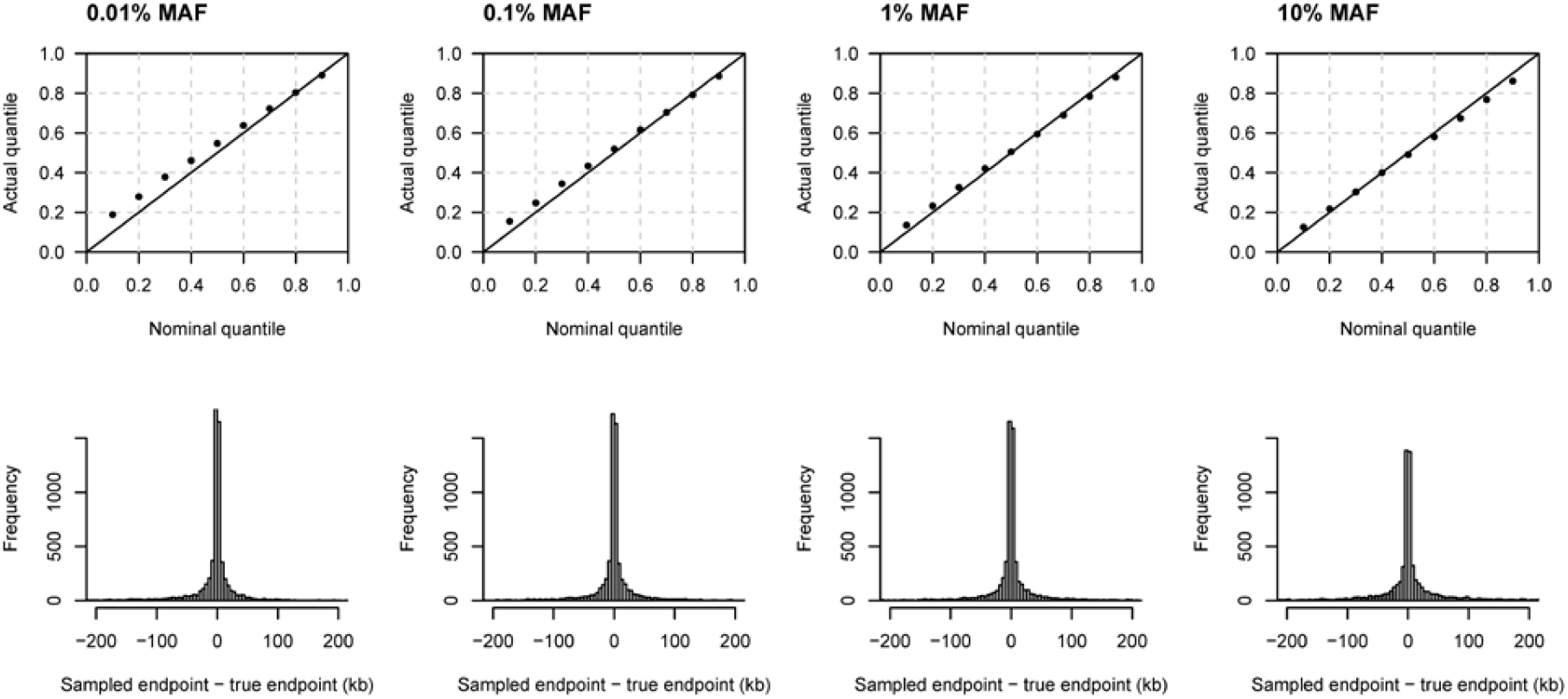
Performance on UK-like simulated data with varying minor allele frequency filters. The data comprise sequence data on 50,000 individuals simulated from a UK-like demographic history (see Methods), with a genotype error rate of 0.02%. The top row shows quantile-quantile plots which assess the calibration of the estimated endpoint uncertainty. The *y* = *x* line is shown for comparison. The actual quantile (y-axis) corresponding to a given nominal quantile (x-axis) is the proportion of segments for which the reported nominal quantile of the right endpoint is greater than the true right endpoint. The bottom row shows histograms of the right endpoint sampled from the estimated posterior distribution minus the true right endpoint of the underlying segment. Histogram bin widths are 5 kb. Results for the left endpoints are similar but are not shown. The true haplotype phase is used in all analyses. The minimum minor allele frequency threshold applied in the ibd-ends analysis is shown above each column. A MAF threshold of 0.01% corresponds to at least 10 copies of the minor allele in these data. Compute times were 4.6 hours (0.01% MAF), 1.4 hours (0.1% MAF), 60 minutes (1% MAF), and 51 minutes (10% MAF). The default MAF for ibd-ends is 0.1%, so the second column shows the same results as the left column of Figure 3.

**Figure S5:**
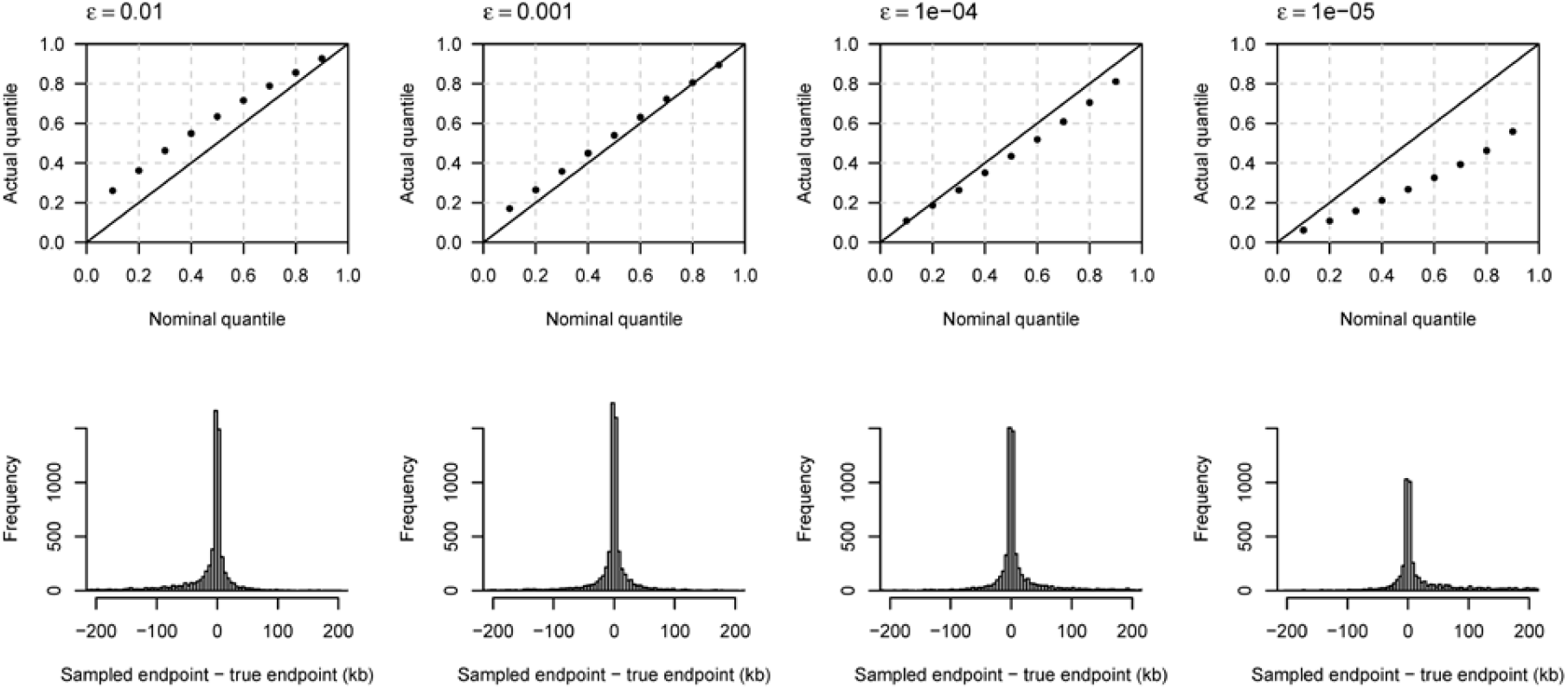
Performance on UK-like simulated data with varying analysis error rates. The data comprise sequence data on 1000 individuals simulated from a UK-like demographic history (see Methods), with a genotype error rate of 0.02%. The top row shows quantile-quantile plots which assess the calibration of the estimated endpoint uncertainty. The *y* = *x* line is shown for comparison. The actual quantile (y-axis) corresponding to a given nominal quantile (x-axis) is the proportion of segments for which the reported nominal quantile of the right endpoint is greater than the true right endpoint. The bottom row shows histograms of the right endpoint sampled from the estimated posterior distribution minus the true right endpoint of the underlying segment. Histogram bin widths are 5 kb. Results for the left endpoints are similar but are not shown. The true haplotype phase is used in all analyses. The analysis error rate, *ϵ*, is shown above each column. The estimated error rates can be found in Table S1.

**Figure S6:**
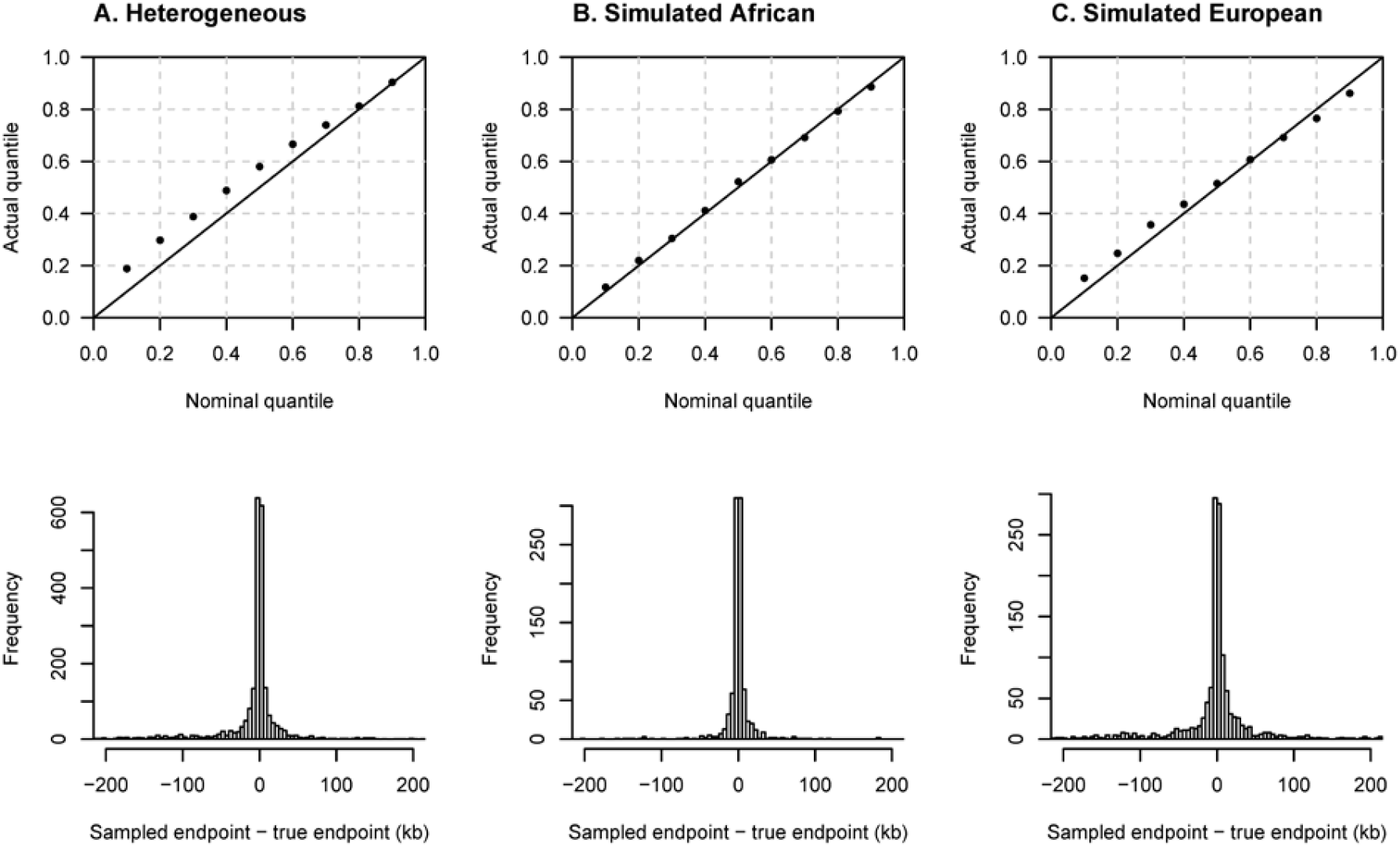
Performance in a simulated two-population model. The data comprise sequence data on 500 individuals each from simulated African and European demographic histories. The top row shows quantile-quantile plots which assess the calibration of the estimated endpoint uncertainty. The *y* = *x* line is shown for comparison. The actual quantile (y-axis) corresponding to a given nominal quantile (x-axis) is the proportion of segments for which the reported nominal quantile of the right endpoint is greater than the true right endpoint. The bottom row shows histograms of the right endpoint sampled from the estimated posterior distribution minus the true right endpoint of the underlying segment. Histogram bin widths are 5 kb. Results for the left endpoints are similar but are not shown. The true haplotype phase is used in all analyses. (A) Combined analysis of individuals of simulated African and simulated European ancestry. (B) Analysis of individuals of simulated African ancestry only. (C) Analysis of individuals of simulated European ancestry only.

**Figure S7:**
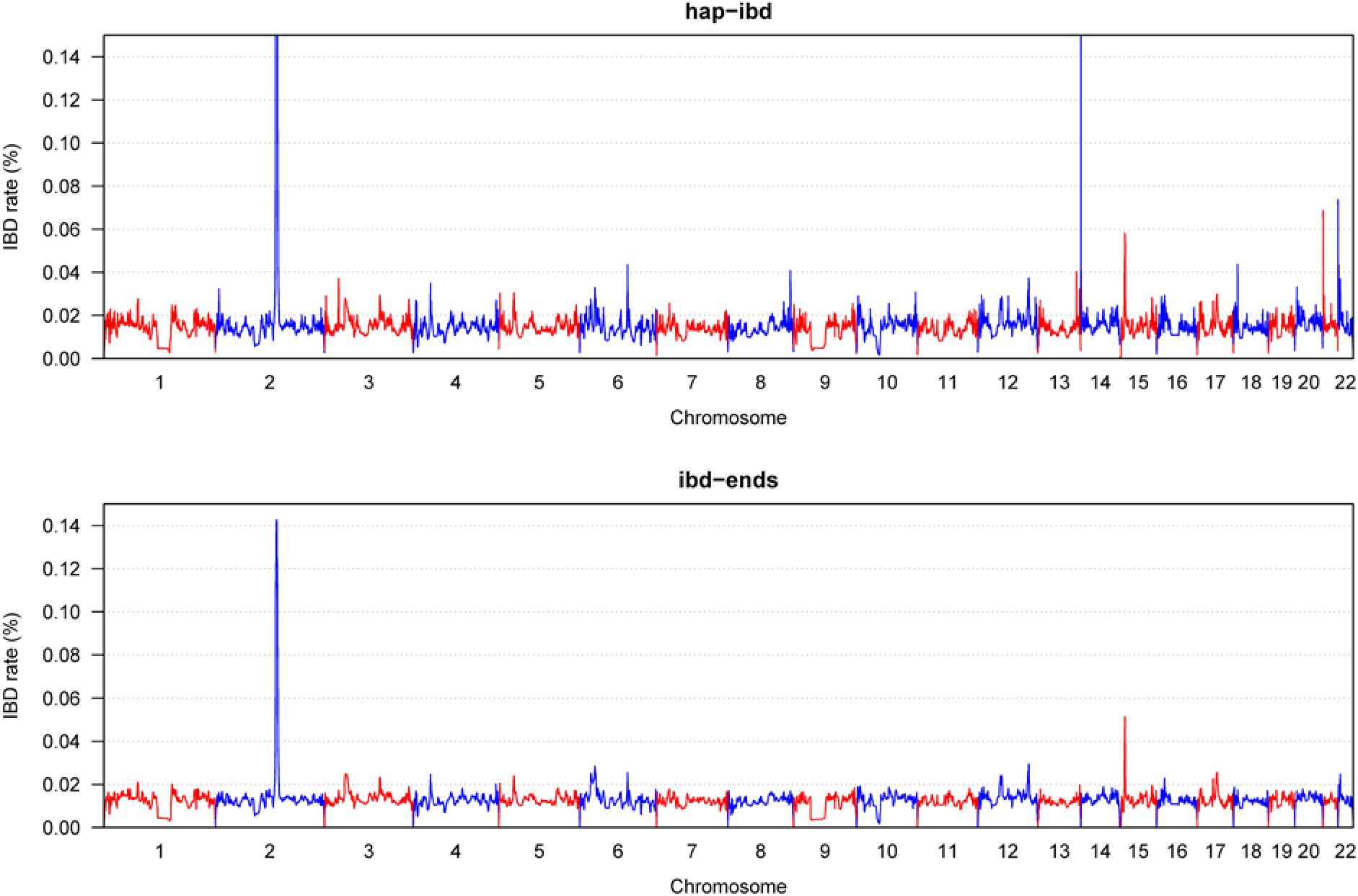
Comparison of rates of IBD detected by hap-ibd and ibd-ends in the UK Biobank White British data. The x-axis shows position along each chromosome. Chromosomes alternate in color. The y-axis is the IBD rate, which is the percentage of pairs of haplotypes for which the position on the chromosome is covered by a sampled IBD segment with length > 2 cM for the haplotype pair. The IBD rate is calculated at 10 kb intervals. The peak heights on chromosomes 2 and 14 for hap-ibd are 0.54% and 2.79% respectively. The lower panel is the same data as in Figure 4.

**Figure S8:**
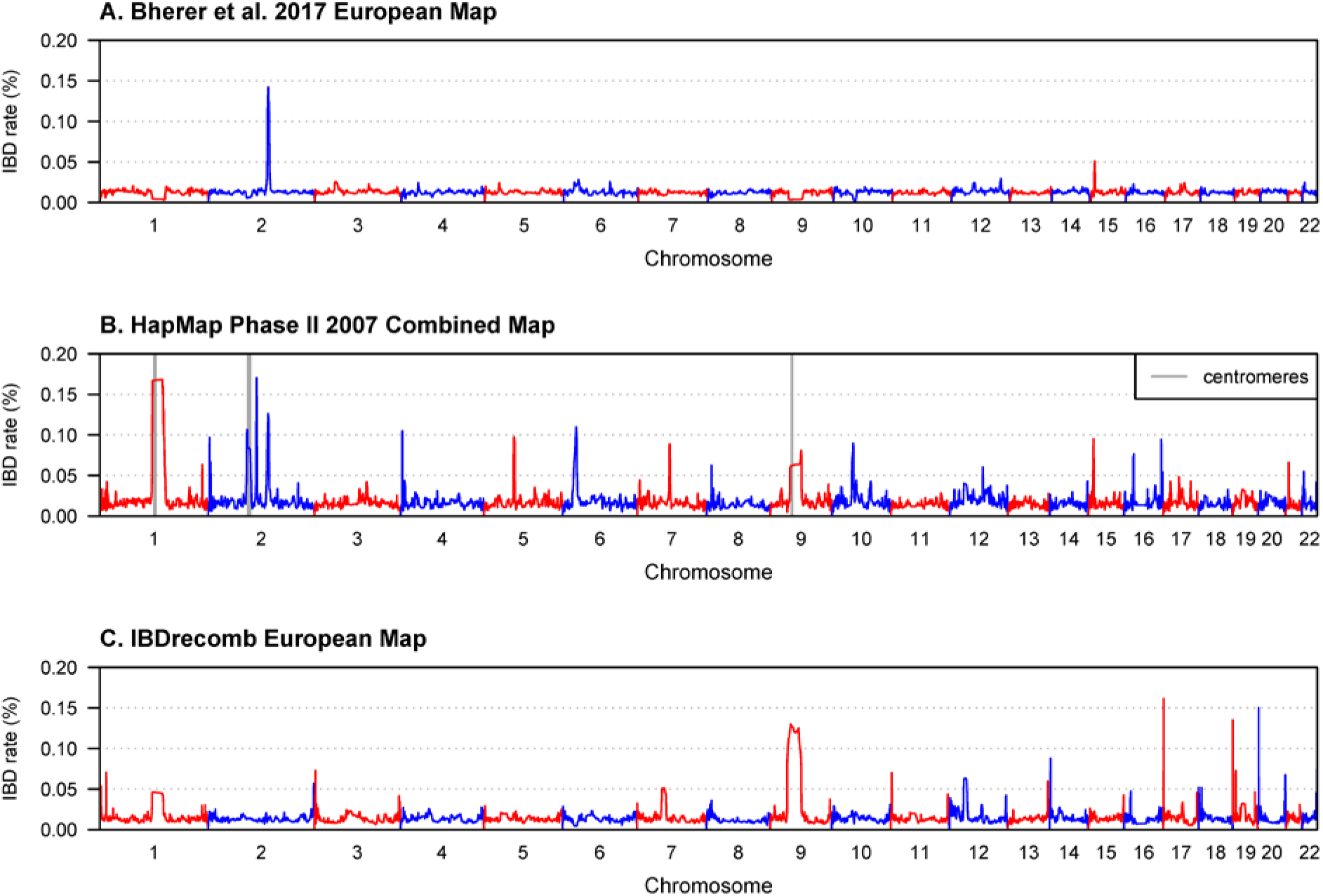
Effect of genetic map on inferred IBD rate. We estimated endpoint uncertainty for 50,000 White British individuals from the UK Biobank using three different maps. A) Bherer et al.’s refined European map.^49^ B) The HapMap Phase II combined ancestry map,^76^ with the centromeres for chromosomes 1, 2, and 9 notated. C) The IBDrecomb European map ^13^ calculated at a 1 kb scale. The y-axis is the IBD rate, which is the percentage of pairs of haplotypes for which the position on the chromosome is covered by a sampled IBD segment with length > 2 cM for the haplotype pair. The IBD rate is calculated at 10 kb intervals.

**Table S1:** Estimated error rates for UK-like simulated data

| **Genotype**<br>**error rate** | **Sample size** | **Sequence or**<br>**SNP data** | **True/inferred**<br>**phase** | **Analysis error**<br>**rate** | **Estimated**<br>**error rate** |
| --- | --- | --- | --- | --- | --- |
| 0.02% | 50,000 | Sequence | True | 0.0005 | 0.00037 |
| 0.02% | 50,000 | Sequence | Inferred | 0.0005 | 0.00032 |
| 0.02% | 50,000 | SNP | Inferred | 0.0005 | 0.00012 |
| 0.02% | 1000 | Sequence | True | 0.0005 | 0.00054 |
| 0.02% | 200 | Sequence | True | 0.0005 | 0.0013 |
| 0.1% | 50,000 | Sequence | True | 0.0005 | 0.0011 |
| 0.1% | 50,000 | Sequence | Inferred | 0.0005 | 0.00096 |
| 0.02% | 1000 | Sequence | True | 0.01 | 0.00065 |
| 0.02% | 1000 | Sequence | True | 0.001 | 0.00057 |
| 0.02% | 1000 | Sequence | True | 0.0001 | 0.00047 |
| 0.02% | 1000 | Sequence | True | 0.00001 | 0.00031 |
| 0.02% | 1000 | Sequence | True | 0.000001 | 0.00012 |

